# Enhancing Breast Cancer Classification with Stacked Convolutional Neural Networks for Whole-Slide Image Analysis

**DOI:** 10.1101/2025.01.02.631156

**Authors:** Tasmiya Mujawar

**Affiliations:** California Lutheran University

## Abstract

This study explores an advanced approach to automatic breast cancer characterization by leveraging convolutional neural networks (CNNs) trained on 224×224 pixel patches from whole-slide images (WSIs). I compared two architectures, VGG-16 and WRN-4-2, across two classification problems: benign versus cancer, and benign, DCIS, versus IDC. Results indicated that WRN-4-2 slightly outperformed VGG-16 on the two-class problem but underperformed on the three-class problem. Stacking CNNs with increased input sizes (512×512, 768×768, and 1024×1024 pixels) significantly improved accuracy, with the 1024×1024 network achieving the highest performance but at a higher computational cost. Dense prediction maps were generated from these stacked networks, enabling effective whole-slide image classification with approximately 90% accuracy for detecting cancer. However, performance for distinguishing between benign, DCIS, and IDC lesions was lower at 76.6%. Future improvements could involve stain standardization, enhanced resolution of prediction maps using architectures like U-net, and comparative studies of multi-resolution approaches to further refine cancer detection and classification accuracy.

## 1 Introduction

Breast cancer is the most prevalent cancer among women globally [16], causing more fatalities than any other cancer type, particularly in developing regions [14]. An essential aspect of breast cancer detection and management is the microscopic analysis of tissue samples by a pathologist.

During this analysis, the pathologist evaluates specific features that indicate the cancer’s potential for growth and metastasis. Together, these features contribute to a semiquantitative assessment of the histologic grade of the disease [5]. The grading deems crucial for predicting prognosis and guiding treatment decisions.

Emergence of *digital pathology* has revolutionized the examination of human tissue by allowing the digitization of slides, facilitating remote consultations among pathology experts. Moreover, digital images enable the application of computer algorithms for automated analysis, enhancing diagnostic capabilities.

Traditional visual interpretation of tissue sections can be tedious and subjective. *Computer-aided diagnosis (CAD)* holds significant promise in mitigating these challenges, thereby reducing the pathologist’s workload. By automating the analysis process, CAD not only provides more precise diagnostic information but also aids clinicians in tailoring optimal treatment strategies for individual patients. Additionally, CAD systems can effectively differentiate benign samples from suspicious ones and quantitatively characterize areas of concern.

### 1.1 Objectives

In this study, I investigate automated outlook for the identifying malignant breast lesions in whole-slide images (WSIs) stained with hematoxylin and eosin (HE). HE staining is the most widely utilized technique in medical diagnostics, routinely employed as part of standard pathological procedures. This staining method imparts a blue color to cell nuclei while rendering the cytoplasm and connective tissue in shades of pink.

The application of HE staining not only facilitates the visual differentiation of cellular components but also provides critical information for diagnosing various pathologies. By automating the analysis of HE-stained WSIs, I aim to enhance the efficiency and accuracy of breast cancer diagnosis, allowing pathologists to focus on more complex cases. Through advanced image processing techniques and algorithms, I seek to improve the identification of malignant lesions, thereby supporting timely and effective treatment decisions for patients.

Previous research on tissue type detection and characterization in pathology images has primarily focused on the categorizaytion. For instance, Bejnordi et al. developed a method for detecting ductal carcinoma in situ (DCIS) at the whole-slide level. Their approach comes with increased distinction of superpixels. From the identified places, handcrafted features are extracted and utilized as feature vectors for statistical models. This method successfully detected 80% of DCIS lesions while maintaining a limit of two false positives per whole-slide image (WSI).

In recent years, *convolutional neural networks* (CNNs) have revitalized previously stagnant fields by achieving high performance in CV and automated speech recognition [1,7]. These deep learning methodologies leverage connectionist models with multiple layers, allowing for the learning of effective feature representations in a supervised manner. This capacity for automatic feature learning often surpasses the performance of traditional handcrafted features [10].

Figure 2 illustrates examples of normal tissue, benign lesions, DCIS, and IDC.

**Fig. 1.**
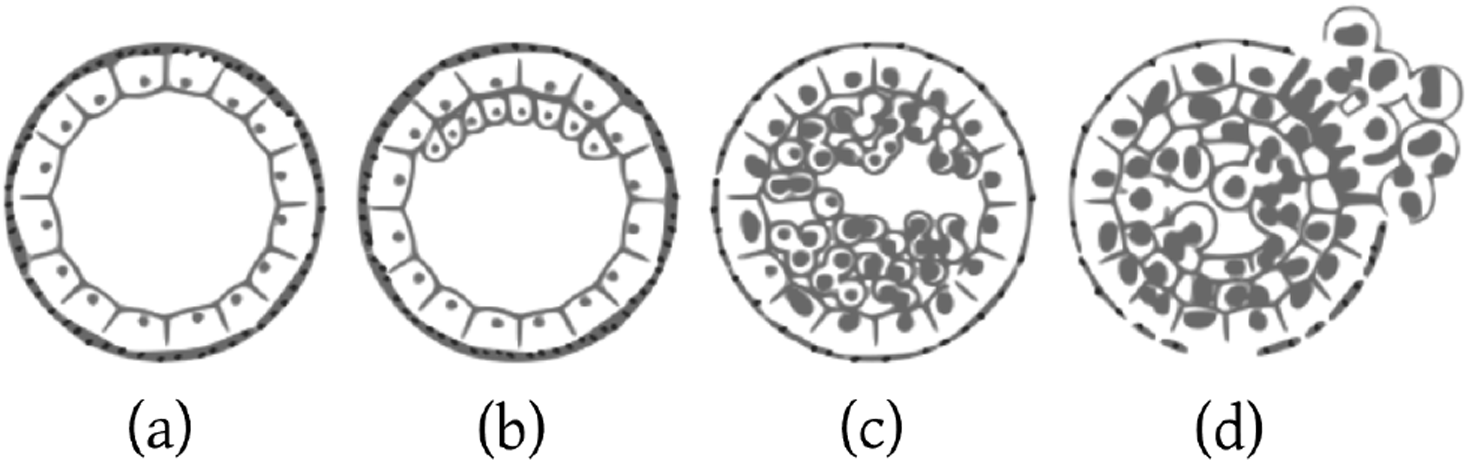
Depiction of mammary ducts.

**Fig. 2.**
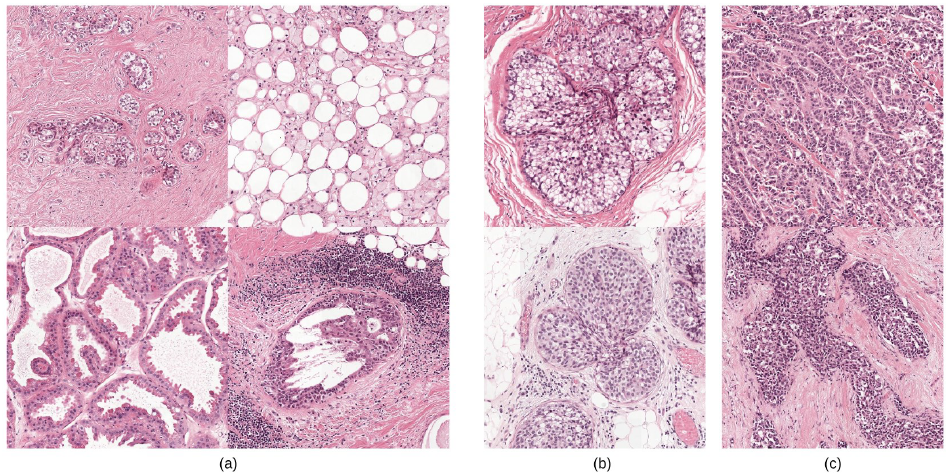
Three classes aimed to identify.

To achieve this automatic labeling, my method comprises several key components. An overview of the complete system is provided in Figure 3. Initially, a fully convolutional neural network (FCN) generates dense predictions for the whole-slide image, where each pixel is assigned a label. Contextual information from the surrounding pixels is crucial for accurate pixel classification. However, due to computational constraints, using large patches with extensive context at high magnifications for training is impractical. To address this challenge, I employ CNN to stop the the result of the previous one.

**Fig. 3.**
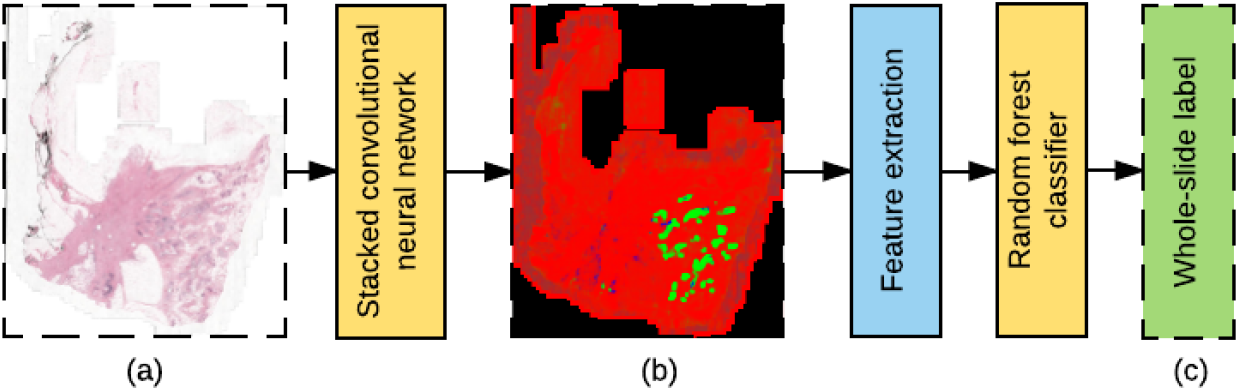
Depiction of labeling system.

I extricate the feature set from the predictions of this stacked CNN architecture, which serves as input for labeling the entire whole-slide image.

For the development and evaluation of the system, I utilize a substantial dataset comprising 221 whole-slide images stained with H&E. This dataset is divided into three distinct sets: one for training classification models, another for intermediate validation and model selection, and a third for final system evaluation (test set). Table 1 provides a breakdown of the number of slides in each set, ensuring no overlap among them.

**Table 1.** Dataset details consisting of 221 whole-slide images.

| LABEL | LESION TYPE | WSI COUNT |  |  |
| --- | --- | --- | --- | --- |
|  |  | *Train* | *Validation* | *Test* |
| *Benign* |  |  |  |  |
|  | *Total* | 50 | 19 | 31 |
| *DCIS* |  |  |  |  |
|  | Grade I DCIS | ? |  |  |
|  | Grade II DCIS | ? |  |  |
|  | Grade III DCIS | ? |  |  |
|  | *Total* | 38 | 11 | 20 |
| *IDC* |  |  |  |  |
|  | *IDC* | 30 | 9 | 13 |
| ***Total*** |  | **118** | **39** | **64** |

## 2 Detailed Classification

Analyzing high-resolution tissue samples, particularly those with magnification levels of up to 40X, presents significant computational and memory constraints for deep learning models, such as CNNs. Whole-slide images (WSIs) are often extremely large, with pixel dimensions reaching 100,000 by 200,000 and uncompressed file sizes nearing 20GB. Each pixel in these images corresponds to an actual area of 0.243 *µm*^2^.

To manage these vast images, a patch-based approach is employed, where smaller sections of the WSI are analyzed individually. The model processes these sections using a sliding window method, gradually producing a comprehensive map of predictions across the entire slide.

Successfully executing this method requires finding a balance between image input size, mini-batch size, and the complexity of the model. Larger image inputs increase memory consumption during training due to the higher number of activations required. Mini-batch size, which dictates the number of images used for each gradient update, can reduce the randomness of the gradient but does not necessarily lead to faster or more accurate learning outcomes [3].

### 2.1 Incorporating Context

Since patches are limited in size, they contain minimal information. While down-sampling larger images to fit patches could result in loss of fine detail, such detail is critical for distinguishing between benign and malignant tissues. Larger scale architectural features become more relevant when differentiating DCIS from IDC.

This method presents images to a multi-stream network at various magnifications. Multiple sub-networks handle different magnification levels, with larger-context patches often downscaled, potentially leading to information loss. Outputs from these sub-networks are fused through fully connected layers, allowing the entire system to be trained via backpropagation.

This paper presents an alternative method for incorporating context while maintaining detail through *stacked CNN*.

### 2.2 CNN

CNNs are composed of various layers, some of which retain spatial information, such as convolutional layers where filters convolve over the image. The filters are not affected by size, allowing images of various sizes to be processed. This property is central to *fully convolutional neural networks*, which excel in segmentation tasks [11].

In my approach, I first train a network on 224 by 224 pixel patches to learn feature representations from fine tissue details. I then remove the bottom surfaces that are unable to meet the criteria, allowing us to increase the patch size (e.g., 768 by 768 pixels) for a forward pass, producing a feature volume resembling an image with multiple feature maps.

The CNN is then trained to optimize memory usage while enabling the processing of high-resolution patches with extensive receptive fields.

### 2.3 Architectures

I evaluate two architectures for patch classification using 224×224 pixel patches. All CNN implementations were developed with Theano and Lasagne frameworks for Python [4, 18], with complete implementations available on GitHub (pending confirmation).

#### Wide Residual Network

The Residual Network won the 2015 ImageNet challenge [8]. Unlike previous architectures, it omits a fully connected layer at the end and can be very deep, with up to 152 layers. It employs residual learning blocks that facilitate learning identity functions when optimal.

I use a variant known as the Wide Residual Network (WRN), as proposed by [19], which shows that wider, shallower networks can outperform deeper, narrower ones in both accuracy and efficiency. My architecture, denoted as *WRN-4-2*, has *N* = 4 (depth) and *k* = 2 (width), balancing batch size and training speed. The ResNet block type is illustrated in Figure 4b. figure 4a depicts the results.

**Fig. 4.**
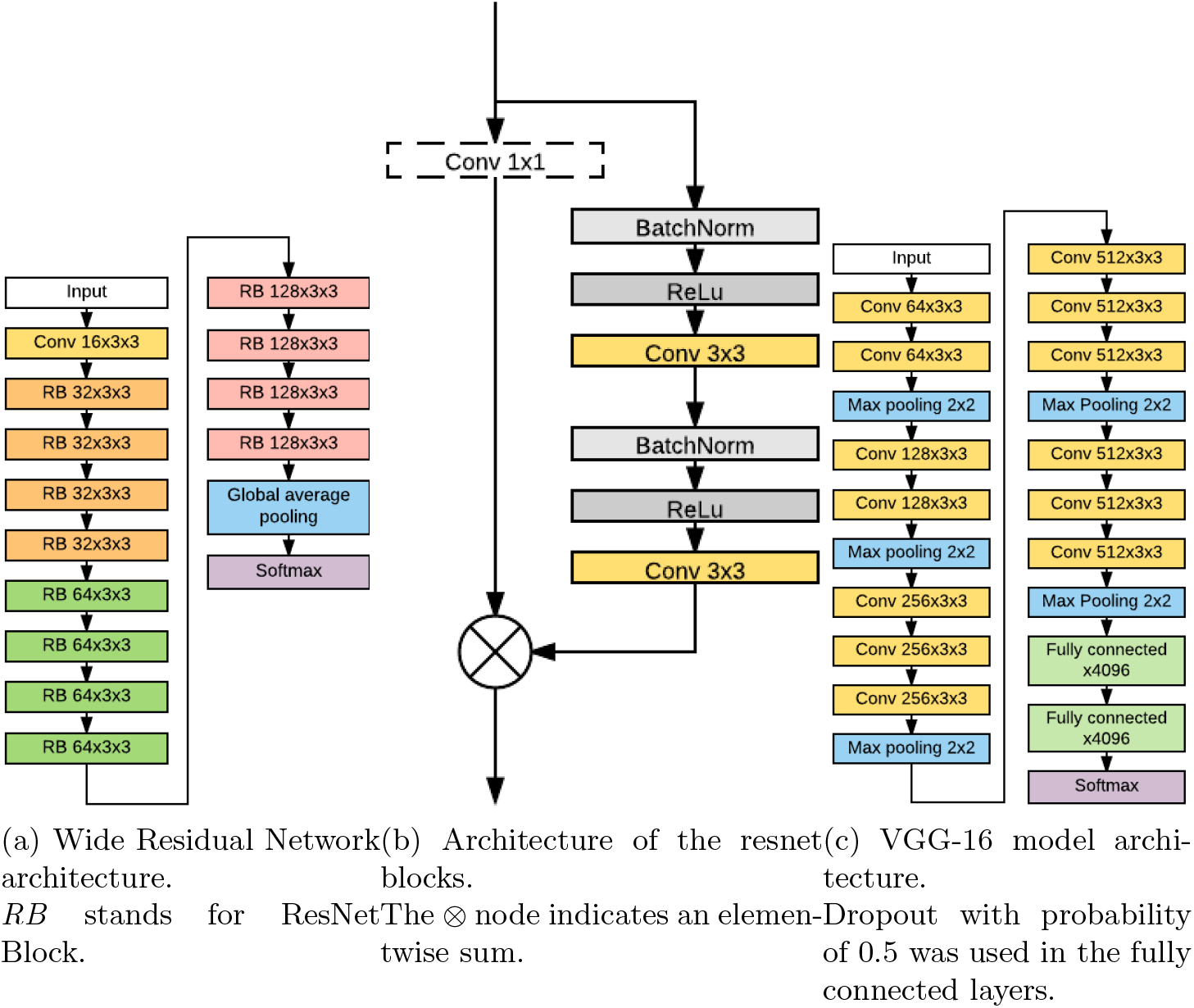
224×224 Wide ResNet downsampling 4a).

**Fig. 5.**
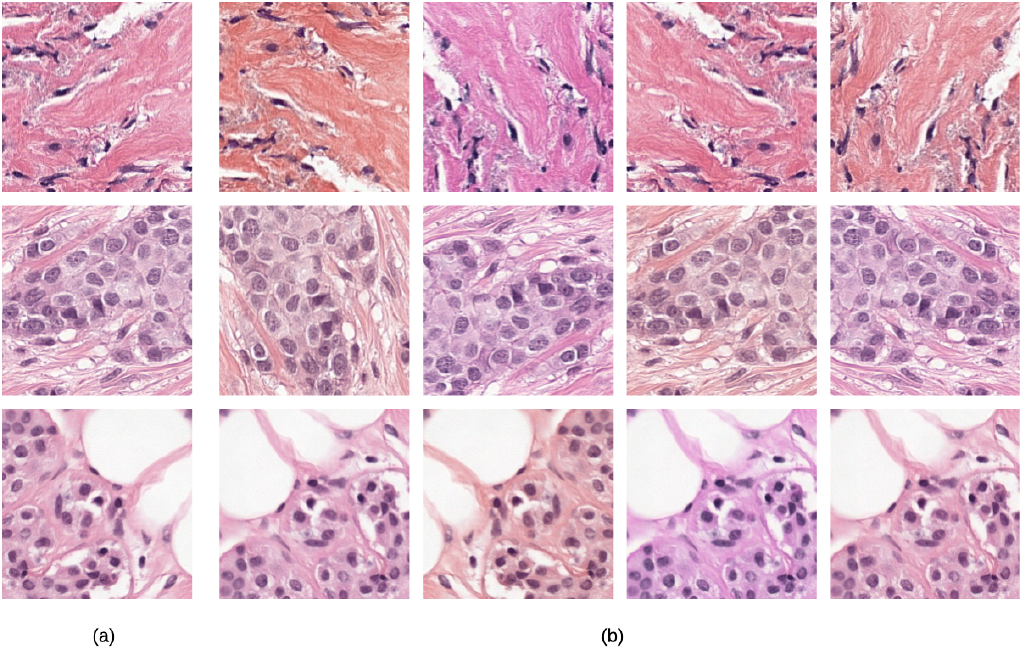
Examples of augmentations applied to 224×224 pixel patches. Note the subtle shifts in hue and saturation in some augmentations.

#### VGG-16

VGG-16 is a widely used framework for image recognition and segmentation, notable for its simplicity [17]. It employs small 3×3 convolution filters and 2×2 pooling across 16 layers. Unlike the ResNet architecture, it lacks skip connections and includes two surfaces at the periphery.

### 2.4 Model parameters

I utilize [12]. The initial learning rates were set to 0.05 for the WRN-4-2 model and 0.01 for VGG-16, as empirically determined to be effective. Both networks employed Rectified Linear Units (ReLU) as nonlinearities, and weights were initialized using He initialization [9]. A mini-batch size of 22 was used for parameter updates, while batches of 128 patches were utilized to assess performance on the validation set. Each patch corresponds to the value of its center pixel in the annotations.

The speed of learning is reduced by 0.2 after 8 stagnant epochs (patience), which increases by 20% after each reduction, while mini-batches are generated by selecting random class samples with uniform probabilities. Table 2 presents the number of annotated regions for each class. One epoch is defined as 300 mini-batches for training and 200 for validation, as the concept of epoch becomes less relevant due to the vast number of pixel locations available for sampling.

**Table 2.** Regions of interest (ROI) per class.

| LABEL | ROI COUNT |  |
| --- | --- | --- |
|  | *Train* | *Validation* |
| Benign | 5097 | 795 |
| DCIS | 1092 | 151 |
| IDC | 2307 | 432 |

Data augmentation enhances the model’s generalization capabilities by introducing variations that do not alter the sample labels. This technique has a regularizing effect that helps prevent overfitting, particularly when realistic variations are applied.

The following augmentation methods are employed:

Zooming is another frequently utilized data augmentation technique; however, it was not incorporated in this study due to the importance of nuclei size in determining class labels. In previous applications of convolutional neural networks, elastic distortions have been successfully implemented as a part of data augmentation strategies, especially in the medical imaging field, including cytology [13] [15]. The variability in nuclei sizes, along with their architectural characteristics, serves as key biomarkers for distinguishing between benign and malignant lesions. Consequently, the use of elastic distortions could potentially obscure this crucial information, leading to its exclusion from my data augmentation approach.

### 2.5 Stacked architecture

I enhance dense predictions by stacking a CNN on top of the WRN-4-2 network’s final layer and compare its performance on binary vs. ternary networks, while evaluating input sizes of 512×512, 768×768, and 1024×1024.

As illustrated in Figure 6, the architecture represents a hybrid model that integrates elements of the Wide ResNet and VGG architectures. This fully convolutional network facilitates faster dense predictions by enabling the reuse of overlapping predictions.

**Fig. 6.**
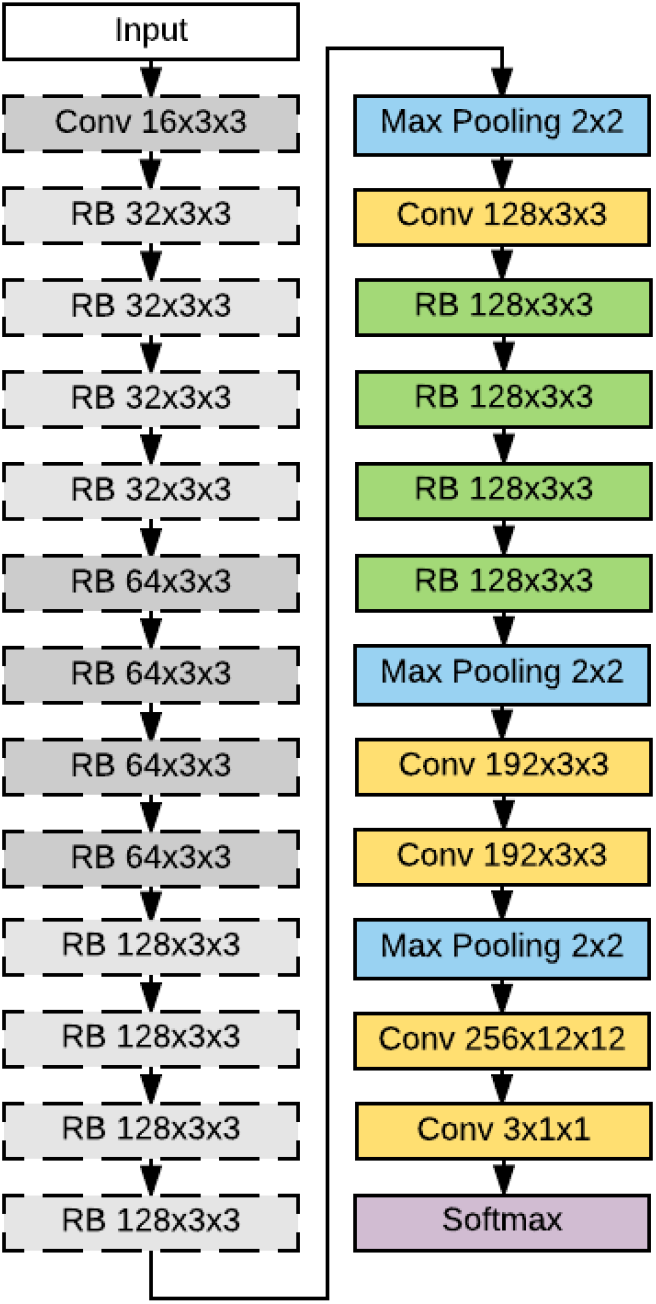
Stacked 768×768 network, frozen.

I train with a learning rate of 0.005 and consistent data augmentations, using minibatch sizes of 18 and 40 for 512×512 and 768×768 patches, respectively, with each epoch consisting of 100 update steps and performance averaged over 75 validation minibatches. Due to the higher memory demands of the 1024×1024 patch network, I limit training minibatches to 10 patches and average validation performance over 125 batches of 22 patches each.

### 2.6 Dense Prediction

Despite the fully convolutional nature of this architecture, I do not increase the input size further. Padding is applied throughout the network, which may introduce learned border effects in the predictions that might not occur with larger input sizes. Although this may not negatively impact performance, I avoid exploiting this option to ensure it does not skew the results. The classifier is shifted by 256 pixels (i.e., a stride of 256) to create a probability map that captures a reasonable probability distribution.

Figure 7 presents examples of the dense predictions for WSIs generated with these parameters, resulting in coarse segmentations of various tissue types.

**Fig. 7.**
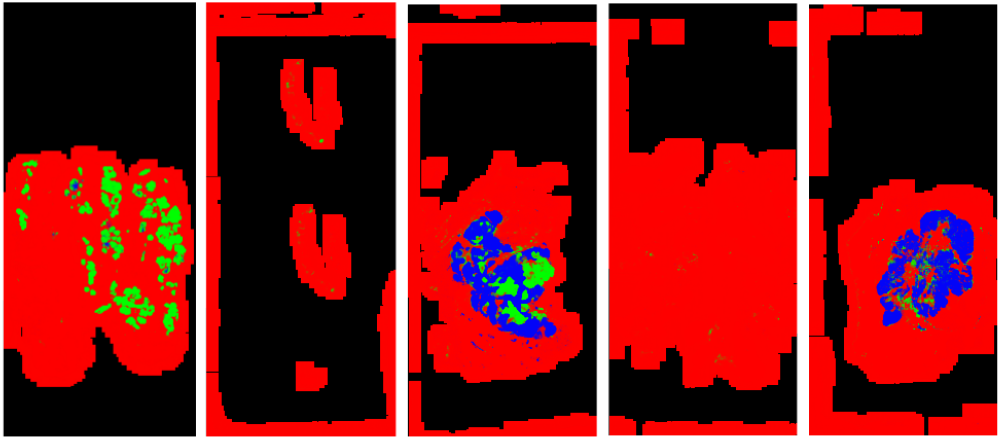
Dense prediction maps: RGB probabilities.

**Fig. 8.**
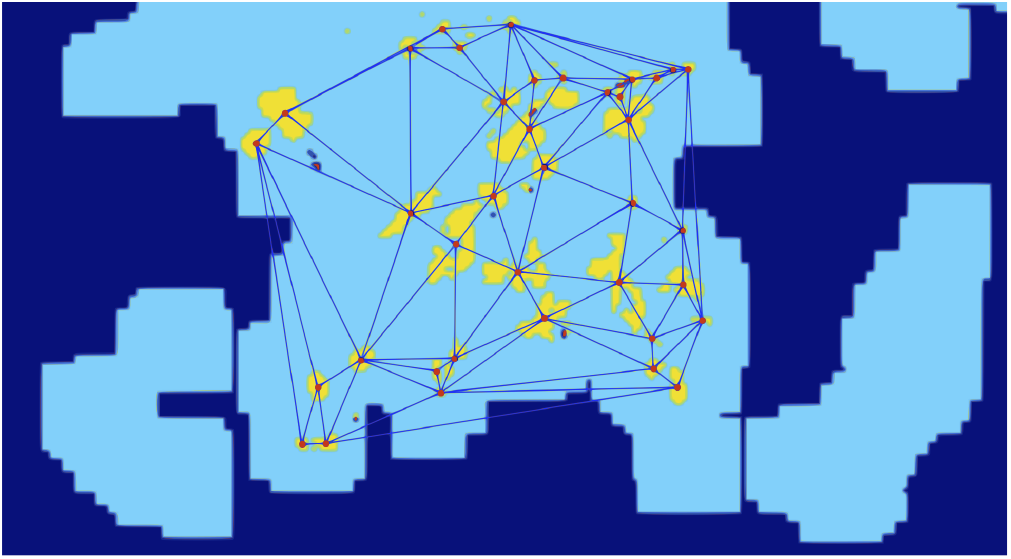
Delaunay triangulation of lesions.

## 3 Whole-Slide Image Labeling

The study aims to classify whole-slide images (WSIs) into benign, ductal carcinoma in situ (DCIS), or invasive ductal carcinoma (IDC) using a dense prediction generated by a stacked CNN. A classifier is trained on features extracted from the resulting probability maps, where each WSI is assigned a single label based on the dominant class. If both DCIS and IDC are present, the image is labeled as IDC.

Three feature sets are extracted:

Full Image Descriptors: Fraction of pixels for each class and their relative proportions among cancerous regions. Voronoi Diagram Features: Metrics derived from Voronoi regions based on connected components, including area, eccentricity, and the number of components. Delaunay Triangulation Features: Metrics from a triangulation graph of connected components, focusing on node distances and the number of neighbors. Convex Hull Area Features: Calculating the fraction of IDC pixels within a convex hull to filter out sparse areas and reduce false positives.

## 4 Feature Vector and Model Training

The feature vector consists of 54 features, which are fed into a random forest model with 512 decision trees, each using a subset of the features (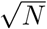, where *N* is the total number). Implemented with Scikit-learn, the model is optimized using leave-one-out cross-validation (LOOCV) due to the limited number of whole-slide images (WSIs). LOOCV involves training on all images except one, then predicting the excluded image, repeating this for each sample. Final evaluation is done on a held-out test set to measure generalizability.

I assess the model using three key metrics:

1. **Accuracy:** Ratio of correct predictions to total predictions, but limited by class imbalance.
2. **Cohen’s Kappa:** Measures agreement between predictions and actual labels, adjusting for random chance, ranging from -1 to 1.

Performance on the training set may appear inflated, which enhances dense predictions for these slides.

### Area Under Curve (AUC)

By adjusting the threshold, I can identify a point for class assignment. I construct a receiver operating characteristic (ROC) curve to illustrate model performance across varying thresholds. The area under this curve represents the probability that a classifier will rank a randomly selected positive instance higher than a randomly selected negative instance [6]. This metric effectively captures the model’s overall performance and its ability to handle uncertainty in labeling.

For my analysis, I opted to calculate AUC only for the binary classification problem distinguishing benign versus cancerous WSIs, given the limited samples available for DCIS and IDC, which could produce an inaccurate and fragmented curve.

## 5 Results

I trained 4 CNN architectures on 224×224 pixel patches, with results in Table 3 showing the best epoch for validation accuracy based on averaged results from two hundred batches of one hundred twenty-eight randomly sampled patches.

**Table 3.** Best epoch patch-level accuracy of 224×224 networks.

| LABELS | ARCHITECTURE | ACCURACY |
| --- | --- | --- |
| *Benign, Cancer* | VGG-16 | 0.9237 |
|  | WRN-4-2 | 0.9241 |
| *Benign, DCIS, IDC* | VGG-16 | 0.8245 |
|  | WRN-4-2 | 0.7995 |

**Table 4.**
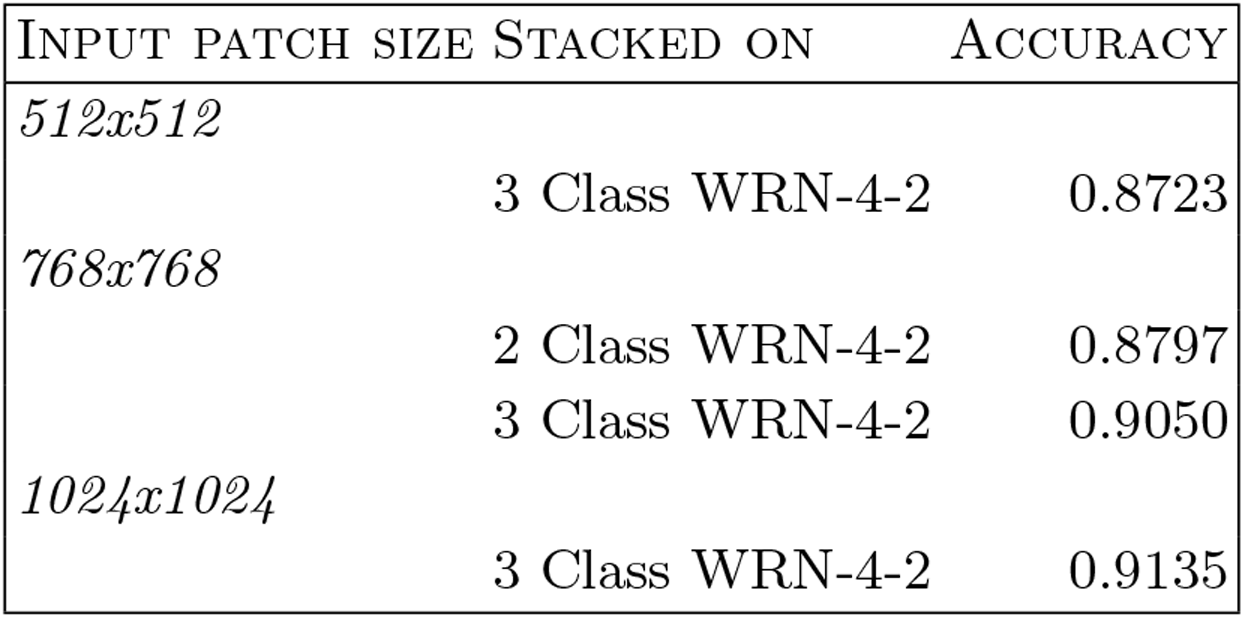
Best epoch patch-level accuracy of stacked networks.

**Table 5.** Confusion matrix of predictions of model used for dense prediction, based on 72,000 random validation set patches.

|  | BENIGN | DCIS | IDC |
| --- | --- | --- | --- |
| BENIGN | 0.878 | 0.088 | 0.034 |
| DCIS | 0.068 | 0.904 | 0.028 |
| IDC | 0.015 | 0.081 | 0.904 |

**Table 6.** Results of whole-slide image label prediction.

| LABELS | RESULT SET | ACC | KAPPA | AUC |
| --- | --- | --- | --- | --- |
| *Benign, Cancer* | Leave-one out CV | 0.8726 | 0.7414 | 0.9264 |
|  | Independent test set | 0.9062 | 0.8123 | 0.9599 |
| *Benign, DCIS, IDC* | Leave-one out CV | 0.8025 | 0.6938 | - |
|  | Independent test set | 0.7656 | 0.6243 | - |

From these results, I can make these observations:

### Architecture Performance

In the binary task of distinguishing benign from cancerous patches, Wide ResNet (WRN) achieved an accuracy of 0.9241, marginally higher than VGG-16’s 0.9237, though not significantly. However, in the three-class task, WRN’s accuracy of 0.7995 fell short of VGG-16’s 0.8245, indicating VGG-16’s superior capacity for distinguishing among the classes. Figure 9 shows WRN’s noisy validation accuracy, likely due to a high learning rate.

**Fig. 9.**
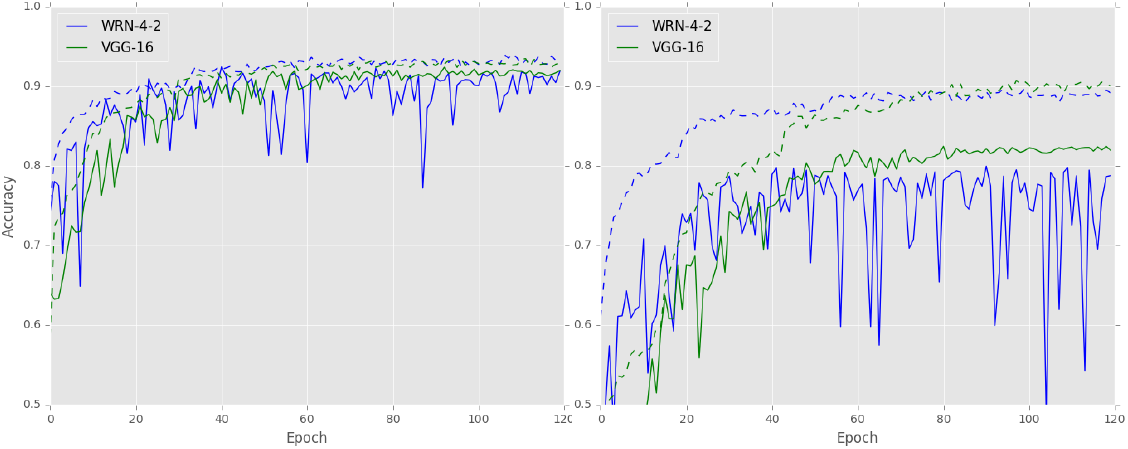
224×224 Network Performance Comparison.

**Fig. 10.**
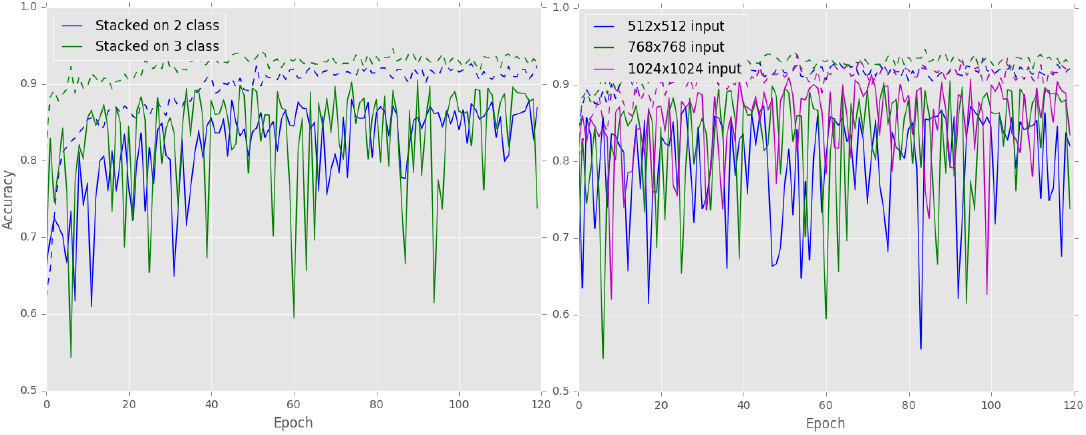
Stacked Network Performance Comparison.

**Fig. 11.**
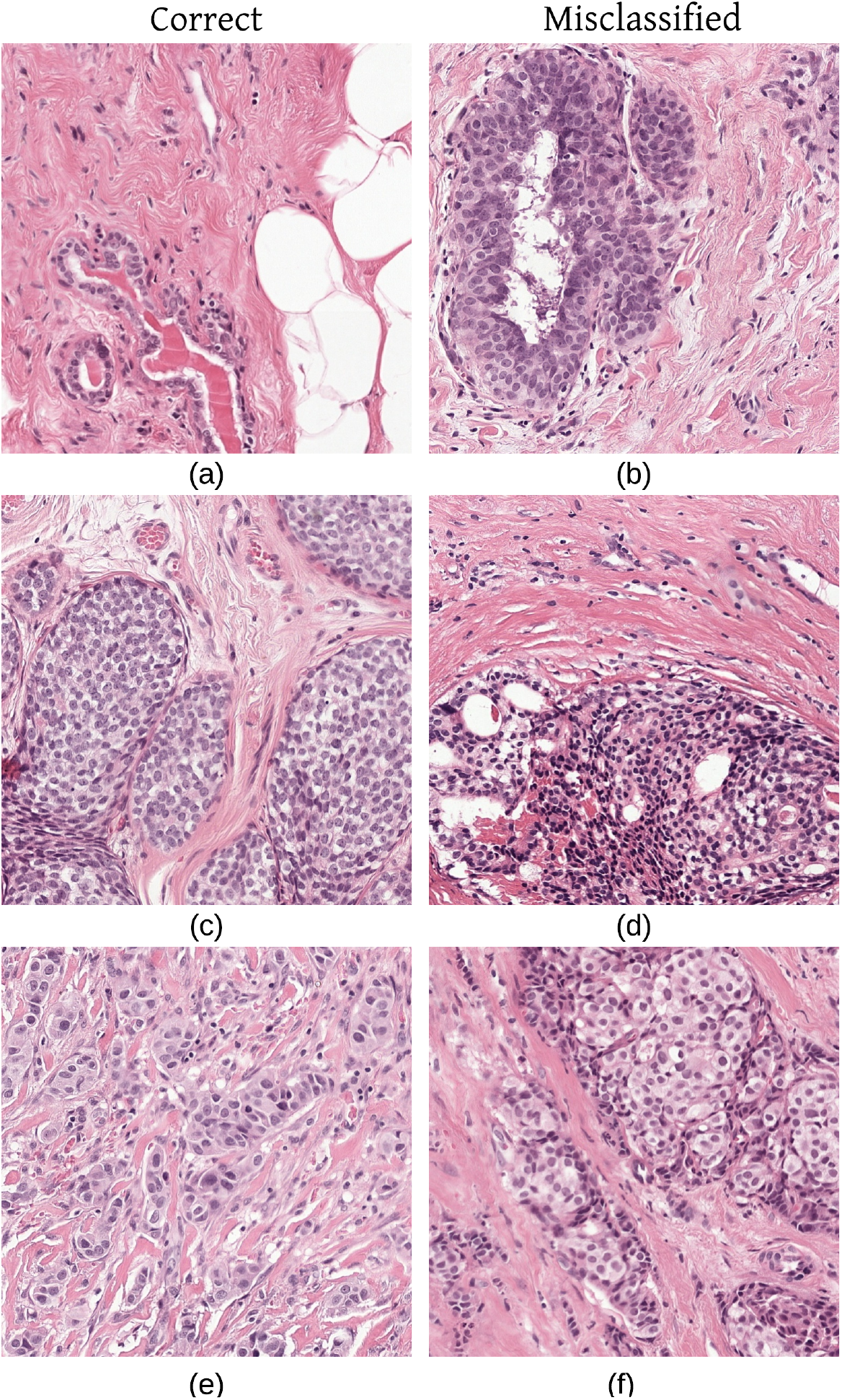
Correct and Misclassified Patches.

The reduced accuracy in the three-class task is linked to limited contextual information in the 224×224 patches, affecting the differentiation between DCIS and IDC. The larger training-validation gap signals overfitting risks. These results are preliminary, as I aim to develop a model that combines outputs from both architectures to enhance performance.

### Architecture Performance

In the binary classification task aimed at differentiating between benign and cancerous patches, the Wide ResNet (WRN) demonstrated superior performance. Despite this slight edge, the difference in accuracy is negligible and may not be statistically significant. However, the dynamics shift in the context of the three-class classification task. In this scenario, the WRN-4-2 network’s accuracy dropped to 0.7995, considerably lower than the VGG-16’s accuracy of 0.8245. This suggests that the VGG-16 architecture is more adept at distinguishing among the three classes being considered. Figure 9 reveals that precision for the WRN is notably erratic, which may indicate that the learning rate was set too high, leading to unstable model updates.

### Two-Class and Three-Class Performance

The observed accuracy for the three-class classification task is significantly lower than that for the binary classification task. This decline can primarily be attributed to the limited contextual information present in the small 224 by 224-pixel patches. Such contextual data is critical for effectively distinguishing between closely related classes, specifically Ductal Carcinoma In Situ (DCIS) and Invasive Ductal Carcinoma (IDC). As depicted in Figure 9, the discrepancy between training and validation accuracies for the three-class networks is much more pronounced. This indicates a heightened tendency for overfitting, which is expected given the insufficient features or context available for the model to learn generalizable patterns.

It is essential to highlight that these results are only preliminary. The individual networks will not be used independently for dense predictions. Instead, I intend to create a higher-level model that incorporates the outputs from both architectures, thereby enhancing predictive performance. This approach aims to capitalize on the strengths of each model, leveraging their unique capabilities to improve overall classification accuracy and robustness in distinguishing between the classes of interest.

The accuracy of all three networks significantly surpasses that of the original models trained on 224×224 pixel patches, indicating a clear improvement in performance. Notably, as the patch size increases, there is a corresponding enhancement in accuracy, suggesting that this improvement is not solely due to the increased depth of the networks. The 1024×1024 model achieved an accuracy of 0.9135, outperforming the smaller architectures, which recorded accuracies of 0.9050 and 0.8723 for the 768×768 and 512×512 networks, respectively, despite the reduced batch size used during training.

However, it is essential to note that the 1024×1024 network is considerably more computationally intensive, both in terms of training time and prediction speed. Given these computational demands, I have opted to utilize the 768×768 network for dense prediction. This model strikes a balance between accuracy and computational efficiency, making it a more practical choice for my application while still delivering robust performance.

The accuracy and kappa coefficient for the held-out test set were significantly lower than the cross-validated results in the three-class problem. Analyzing the class imbalance, the confusion matrix in Table 7 indicates that misclassifications between the DCIS and IDC classes are particularly prevalent, contributing to the disparity in performance metrics.

**Table 7.**
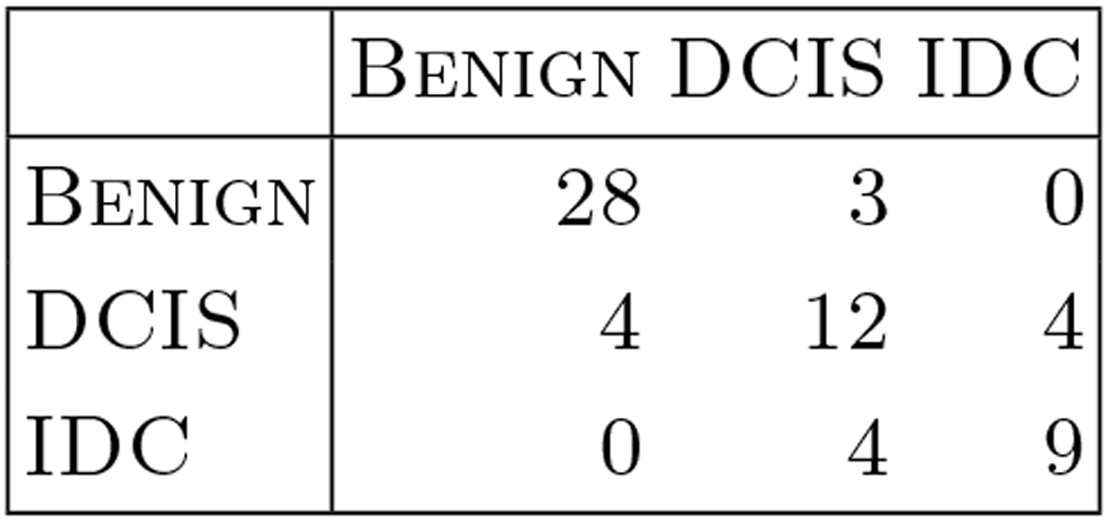
Confusion matrix of test set predictions.

|  | BENIGN | DCIS | IDC |
| --- | --- | --- | --- |
| BENIGN | 28 | 3 | 0 |
| DCIS | 4 | 12 | 4 |
| IDC | 0 | 4 | 9 |

Interestingly, the combined training and validation sets was 0.9264, while the independent test set achieved an AUC of 0.9599. With zero false positives, I anticipate that approximately half of the slides can be confidently labeled as cancerous (see Figure 12).

**Fig. 12.**
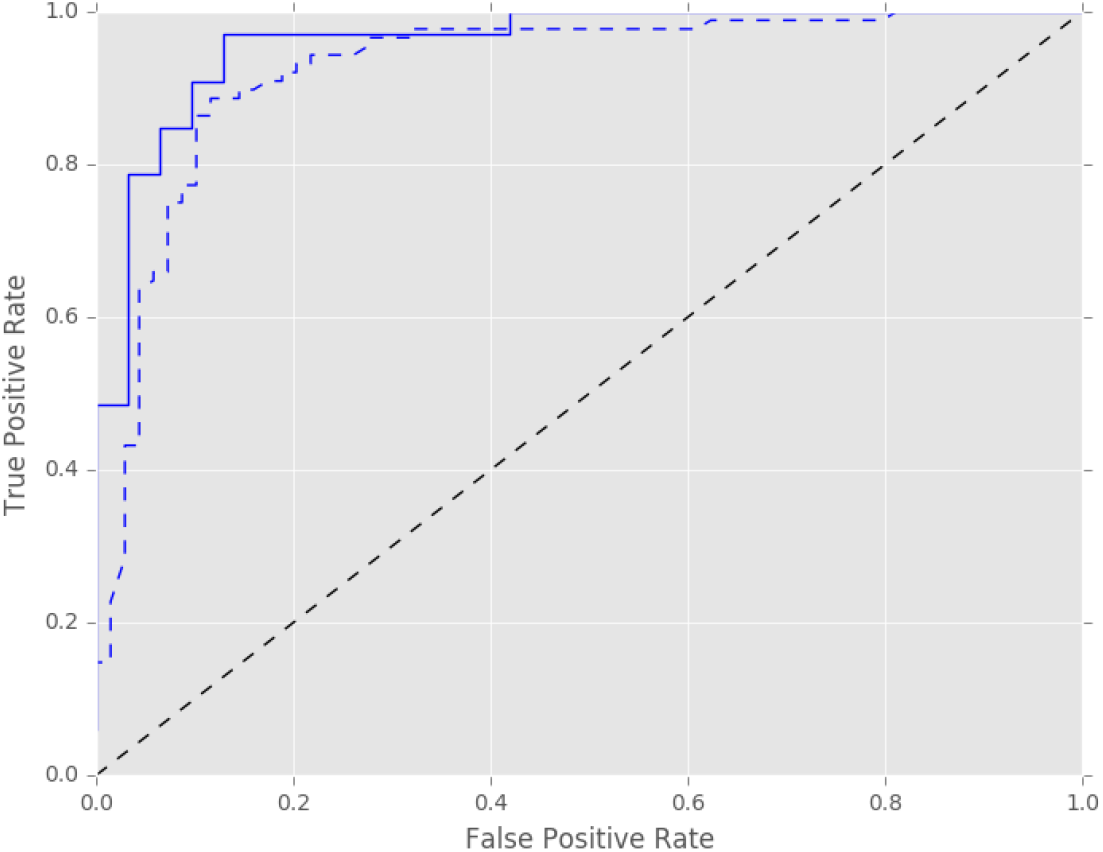
Receiver operator characteristic (ROC) curve. The solid and dashed line are for the independent test set and cross validated train set respectively.

## 6 Conclusion

I assessed an automated method for characterizing breast malignant lesions in whole-slide images (WSIs) stained with hematoxylin and eosin (H&E). By stacking a convolutional neural network (CNN) with a larger input size atop a pre-trained network, I enhanced the context available for predicting image patch labels. This approach yielded significantly better performance compared to smaller networks.

Using these stacked CNNs, I performed a rough segmentation of cancerous areas in the WSIs. From the resulting probability maps, I achieved approximately 90% accuracy in predicting whether a slide contained cancer. On a small independent test set, my system could effectively filter out around 50% of obviously benign slides without error.

However, the accuracy for the three-class problem—distinguishing between benign, Ductal Carcinoma In Situ (DCIS), and Invasive Ductal Carcinoma (IDC)—was lower, at about 76.6%. I anticipate that improvements in dense prediction will enhance whole-slide image classification. With highly accurate dense predictions, labeling entire WSIs would become straightforward. Potential avenues for improvement include:

1. Stain Standardization: Although the current system achieves some hue and saturation invariance, implementing specialized stain standardization techniques could further enhance performance, as demonstrated in previous studies [2].
2. Resolution and Precision of Probability Maps: The existing prediction map is coarse due to the patch-based approach and high stride. Employing fully convolutional network architectures like U-Net [15] could yield finer segmentations, allowing for the extraction of more detailed features that describe lesion morphology.
3. Architectural Comparisons: Future research could explore the effectiveness of my context-enhancing method—stacking CNNs—against multi-resolution architectural approaches.

